# Molecular determinants of low affinity complexes formed by the electrical synapse proteins Connexin 36 and ZO-1

**DOI:** 10.1101/2025.10.27.684941

**Authors:** Stephan Tetenborg, Fatemeh Ariakia, Nikki Brantley, Georg Zoidl, Eyad Shihabeddin, Christophe P. Ribelayga, Tarsis G. Ferreira, John O’Brien

## Abstract

Although chemical and electrical synapses function fundamentally differently, they evidently share common design principles. Like neurotransmitter receptors, Connexin 36 (Cx36) containing gap junction channels, key constituents of electrical synapses, are anchored to scaffolding proteins that stabilize the connexin at the synapse. One of the most prominent proteins that has been described in this context is the Zonula occludens protein 1 (ZO-1). ZO-1 interacts with Cx36 via one of its three PDZ domains. This interaction is inherently weak and was suggested to facilitate the dynamic regulation of electrical synapses. In the present study, we have combined Gaussian accelerated molecular dynamics simulations and binding assays to identify the exact residues in the PDZ binding motif of Cx36 that are necessary to sustain these low affinity interactions. Among the different Cx36 mutations we have generated, we discovered a single substitution at position 319 within the PDZ binding motif that massively increases binding in different experimental settings. In addition to this site, we found that acidic residues adjacent to the PDZ binding motif (PBM) in Cx36 and its fish orthologues are evolutionarily tuned to weaken PDZ interactions as well. We were able to enhance PDZ1 binding drastically by substituting these residues with hydrophobic or positively charged amino acids. Finally, we demonstrate that the weak PDZ1/Cx36 interaction is sensitive to CaMKII mediated phosphorylation of Cx36, suggesting that ZO-1 unbinding may be a necessary event to potentiate electrical synapses. In summary, our study provides a detailed analysis of different mechanisms that can be exploited to modify the interaction between two key components of an electrical synapse: Cx36 and ZO-1.

## Introduction

Neurons have the remarkable ability to communicate via hundreds to thousands of synapses, which serve as the most fundamental information processing units in the nervous system. A synapse is not a static entity but subject to constant changes that alter the strength of transmission in response to neuronal activity, a feature that is known as synaptic plasticity (3, 4). For chemical synapses, plasticity is often described in the context of long-term potentiation (LTP), a well described mechanism that enhances synaptic transmission after periods of frequent stimulation. At the molecular level, LTP is achieved by the increased insertion of AMPA receptors into the postsynaptic membrane, which ultimately increases the excitability of a synapse (5). In order for a synapse to be flexible and grow it is critical that its receptors are anchored to scaffolding proteins residing in the postsynaptic density such as PSD95 (postsynaptic density 95). The capacity of these proteins to cluster receptors and ion channels on the cell surface is not only important for synaptic plasticity but also determines the formation of chemical and electrical synapses during the development of the nervous system (6-9).

A characteristic feature of scaffold proteins is the use of multiple protein-protein interaction domains that mediate multivalent interactions to bring synaptic proteins into close proximity and organize functional signaling complexes into nano domains (10). Although the role of scaffolds of chemical synapses has been studied in great detail, we currently lack a comprehensive understanding of the electrical synapse equivalents of these proteins (11, 12), which is owed to the fact that chemical synapses are predominant in the nervous system. Despite their lesser stake, electrical synapses are widely recognized as essential elements of neuronal circuits given their speed of transmission and distinct functions within circuits. Unlike chemical synapses, electrical synapses do not rely on neurotransmitters for communication, but they instead transmit neural signals instantaneously as ionic currents via intercellular gap junction channels. This seemingly simple form of transmission facilitates the synchronization of neuron populations and thereby regulates complex brain functions, such as vision (13, 14) and the coordination of motoric functions or hormone secretion (15, 16). In the nervous system, electrical synapses are mainly, but not exclusively, formed by Connexin 36, a gap junction protein that is primarily expressed in neurons and often referred to as “the major neuronal connexin.” Gap junction channels that are made of this particular isoform were shown to be highly modifiable and are regulated via phosphorylation, which allows neurons to adapt the extent of electrical coupling to environmental conditions (17-21). For instance, the coupling of AII amacrine cells in the mammalian retina is low under bright daylight due to the release of dopamine, which closes gap junction channels via Protein phosphatase 2A (PP2A) mediated dephosphorylation of Cx36 to increase the spatial resolution of the visual system (17, 18, 22). Under dim light conditions, when it is beneficial to operate with high sensitivity, AII cells become increasingly coupled. This increase in coupling is driven by the presynaptic glutamate release from ON bipolar cells inducing the activation of CaMKII, which in turn phosphorylates Cx36 and opens up the channel. These activity-driven changes in electrical coupling resemble the type of plasticity we have outlined earlier for chemical synapses. In addition to immediately acting mechanisms such as phosphorylation, which occur on a minutes time scale, Cx36 channels exhibit fast turnover rates and short halftimes of 3 ½ hours (23, 24). This implies that an electrical synapse constantly has to replace its channels within a short amount of time and maintain a balance between channel insertion and internalization to remain functional (24). The dynamic nature of these processes raises the question of what kind of mechanisms electrical synapses have evolved to accommodate these constant changes?

Like neurotransmitter receptors, Cx36 channels are embedded into a synaptic density, harboring signaling and scaffold proteins that function as a fundament on which the electrical synapse is built up (2, 6, 7, 12, 25). One of the most prominent interactors that has been described in this context is the Zonula occludens protein 1 (ZO-1). This scaffold was initially identified as a component of tight junctions (26). Later studies demonstrated its association with multiple connexins, including Cx36 (27-30). ZO-1 binds connexins or tight junctions proteins such as claudins with one of its three PDZ domains, small globular domains interacting with short sequences that are presented at the C-terminus of specific binding partners (31). Besides its three N-terminal PDZ domains, ZO-1 also contains an SH3 domain and an actin binding region in its C-terminal portion that is necessary to tether the junctional complex to the cytoskeleton. Genetic screens have shown that certain ZO-1 variants in zebrafish are essential for electrical synapse formation (6, 32), which makes it difficult to assess how these proteins could potentially impact dynamic processes such as phosphorylation or turnover of intact Cx35 channels. Interestingly, surface plasmon resonance measurements (SPR) have demonstrated that the binding interaction of Cx35 (the fish orthologue of Cx36, identical PDZ binding motif) and the first PDZ domain of ZO-1 is transient and inherently weak, with K_D_ values of around 60.3 µM. In contrast, Cx43 and the second PDZ domain of ZO-1 were able to form far more stable complexes with K_D_ of 15.5 µM in the same experimental paradigm (33). Based on these observations, a more regulatory role for ZO-1 has been proposed. One possibility is that the Cx35/ZO-1 interaction is evolutionarily tuned to be weak and transient in order to facilitate fast changes in electrical coupling. How this weak interaction is achieved and how it is affected by posttranslational modifications such as phosphorylation is still unknown.

In the present study, we have combined binding assays and Gaussian accelerated molecular dynamics simulations (GaMD) to identify amino acids in the PDZ binding motif of Cx36 that determine low affinity interactions with ZO-1. By comparing PDZ1 binding for a series of Cx36 mutants we uncovered critical sites that, when mutated, cause drastic changes in ZO-1 binding. In addition to these effects, we further demonstrate that the Cx36/ZO-1 interaction can be regulated via phosphorylation of Serine 315, which provides a mechanistic explanation for the potentiating effects of this site. In summary, our study offers a detailed analysis of the molecular determinants the Cx36 protein has evolved in order to sustain low affinity interactions with ZO-1.

## Results

### The A319T substitution in Cx36 massively increases ZO-1 binding

The binding interaction of Cx36 and the first PDZ domain of ZO-1 is inherently weak, which has been suggested to allow the formation of transient but more dynamically regulated protein complexes that facilitate fast changes in electrical coupling as described for various circuits (18, 19, 21, 33, 34). In a previous study, we have demonstrated that this interaction can be massively increased by an internal deletion in the C-terminal that brings upstream residues RT adjacent to the terminal YV to change the wild type PDZ binding motif SAYV to RTYV (referred to as Δ313-319 in sequence alignment, figure 1B) (1). Given the strong effect on binding, which represents the polar opposite of wild type Cx36, we reasoned that the RTYV mutant could serve as a useful tool to understand the functional relevance of weak PDZ interactions for the regulation of electrical synapses. To identify the amino acids responsible for the shift in binding we substituted the two initial amino acids in the PBM of Cx36 separately with the corresponding residues of the RTYV mutant (SA318-319RT: Serine 318 and alanine 319 to arginine and threonine respectively; S318R: Serine 318 to arginine and A319T: alanine 319 to threonine, all mutations are illustrated in the sequence alignment, Figure 1B) and tested their ability to bind Venus-PDZ1 in CO-IP (GFP-Trap) experiments from cell lysates of co-transfected HEK293T cells. Consistent with our previous reports (1), we detected a massive increase in PDZ1 binding for the Δ313-319 mutant in comparison to wildtype Cx36 (Figure 1C, blue asterisk indicates faint dimer bands for wild type Cx36, red asterisk indicates Cx36 dimers for mutants and the short arrows monomers). We were able to replicate this binding shift with the SA318-319RT and A319T mutant respectively, indicating that the presence of threonine at position 319 alone causes the increase in PDZ binding we have described earlier (1) (Figure 1C).

**Figure 1:**
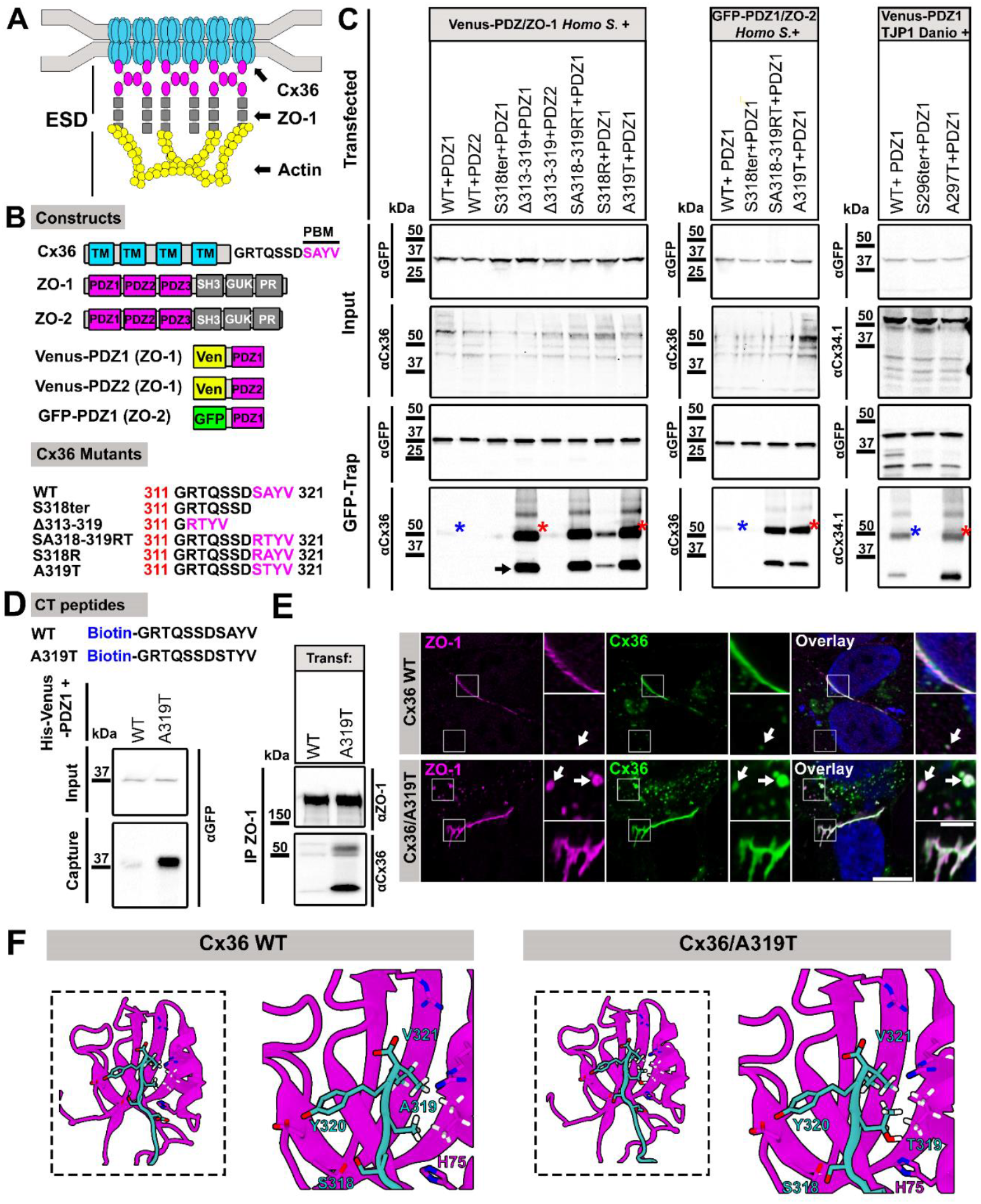
Substituting alanine 319 with threonine in Cx36 massively increases ZO-1 binding. **(A)** Cartoon illustrating the interaction of Cx36 and ZO-1 at electrical synapses. **(B)** Domain organization of constructs used in this study. **(C)** GFP-Trap pull-down of Venus PDZ1/ZO-1 and different Cx36 C-terminal mutants co-expressed in HEK293T cells. The massive increase in PDZ1 binding previously described for the Cx36Δ313-319 (1, 2) can be replicated with a single substitution of alanine 319 to threonine. **(D)** Comparison of His-Venus-PDZ1 binding of C-terminal peptides comprising the amino acid sequence of wildtype Cx36 and the A319T sequence. Consistent with our IP data, the A319T peptide shows a massive increase in PDZ1 binding, whereas binding of the WT peptide is relatively weak. An increase, although more moderate, is observed when A319T binding is tested with the first PDZ domain of ZO-2. The interaction of Cx36-A319T with Zebrafish Tjp1b (ZO1b) PDZ1 was also increased. **(E)** Co-IP experiment of endogenous ZO-1 and the transfected A319T Mutant from HeLa cell lysates. A319T but not wildtype Cx36 is co-precipitated. Confocal scans of both Cx36 variants and endogenous ZO-1 in HeLa cells. ZO-1 (Magenta) is concentrated at gap junctions. Scale: 10 µm. Magnified inset: 2.5 µm. **(F)** Model of the PBM of WT Cx36 and the A319T mutant binding the first PDZ domain of ZO-1.

In addition to PDZ1, we also tested the interaction of the RTYV mutant and the second PDZ domain of ZO-1. This domain was shown to be incompatible with the PBM of Cx36 (27) and therefore served as a negative control to exclude potential changes in PDZ domain specificity that may result from the A319T substitution. In these experiments, however, we observed only little binding, indicating that the preference for PDZ1 is retained.

Next, we asked if the A319T mutant also enhances the binding interaction with PDZ domains of other known interactors of Cx36. ZO-2 is another prominent Cx36 interactor (2, 35, 36). The A319T mutation caused a substantial but slightly smaller increase in binding, while elimination of the PBM (S318ter) eliminated binding (Figure 1C). Finally, we tested whether the principle of PDZ domain binding enhancement by the T mutation is evolutionarily conserved by introducing the equivalent mutation (A297T) into the identical C-terminus of zebrafish Cx34.1. Binding to PDZ1 domain of zebrafish Tjp1b (ZO1b) was also increased, although not to the degree seen with human ZO1 (Figure 1C).

To back up our findings with an additional approach, we performed pull-down assays with synthesized peptides containing an N-terminal biotin and the 11 C-terminal amino acids of wild type Cx36 and the A319T mutant. Each peptide was coated onto streptavidin beads and probed with purified His-Venus-PDZ1 (ZO-1) to assess binding. In line with our CO-IP data, we observed a massive increase in binding for the A319T mutant (Figure 1D). To further assess if the threonine substitution could potentially serve as a tool to boost PDZ interactions in an *in vivo* system, we transfected the A319T clone into HeLa cells and tested if the increased affinity for PDZ domains is sufficient to attract the endogenous pool of ZO-1. In IP experiments with monoclonal ZO-1 antibodies, we observed substantial binding of the A319T mutant but no signs of wildtype Cx36. (Figure 1E). Similar effects are seen in fixed HeLa cells, stained for Cx36 and ZO-1. Although we observed concentrated ZO-1 labeling at gap junctions formed by each Cx36 variant (magnified square), we noticed that only the A319T mutant colocalized with cytoplasmic ZO-1 clusters (arrows). Hence, the increase in binding we have seen in IP experiments most likely arises from these cytoplasmic interactions, because most of the connexin that is produced in transiently transfected HeLa cells accumulates as non-junctional protein in intracellular vesicles (37, 38) (applies to every cell type we have tested). These results confirm that the shift in affinity for the A319T mutant is sufficient to attract substantial amounts of endogenous ZO-1.

To understand the stereochemistry underlying the increase in binding, we combined structural modeling and GaMD simulations. This strategy allowed us to compare to what extent non-covalent interactions of both Cx36 variants with PDZ1 (ZO-1) differ. Figure 1F illustrates a model of PDZ1 binding the PBM of both Cx36 variants based on the PDB structure 2H2B (39). The interaction involves several hydrogen bonds that are formed by backbone and sidechain atoms of the peptide with residues within the PDZ1 groove. In the WT, these hydrogen bonds are particularly enriched at the C-terminal motif, where residues A24, I25, and S26 provide a stable network of contacts with PDZ1. This is reinforced by adjacent polar interactions involving G27, D30, and N31, which stabilize the binding orientation through repeated hydrogen bonds and water-mediated bridges. Additional contributions come from P32, H33, and F34, which engage in hydrophobic and π-stacking interactions, while the conserved S44 and D45 residues in PDZ1 establish recurrent electrostatic contacts. Importantly, distal contacts such as those involving E58, H75, V79, and R83 expand the interaction surface, ensuring that the WT peptide maintains a broad and persistent interface with PDZ1. These networks might explain why the WT PBM engages in a multivalent interaction mode with hydrogen bonds at the peptide core coupled with long-range electrostatic and hydrophobic contacts that extend binding beyond the canonical carboxylate recognition site.

The free energy landscapes of these interactions are illustrated as heat maps (Figure 2A, B) in which areas with the lowest free energy (deep blue areas, outlined) represent the bound state of each amino acid. GaMD simulation of the wildtype PBM and PDZ1 revealed that alanine 319 (Cx36) is rather mobile, occupying distances from 3-11 Å from histidine 75 (ZO-1), while tyrosine 320 (Cx36) remains about 3.5 Å from aspartate 45 (ZO-1)(Figure 2A). In contrast, threonine 319 of the A319T mutant adopts a single stable position about 3.5 Å from histidine 75, while tyrosine 320 remains about 7.5 Å from aspartate (Figure 2B). Ribbon diagrams of these interactions in an unbound and bound state are illustrated for WT (Figure 2A’) and A319T (Figure 2B’). These simulations expose a clear difference in the stability of interactions that are formed by both amino acids: Alanine (WT) is mobile and unstable whereas threonine (A319T) adopts a stable position, which could explain the increase in binding we have seen in pull-down experiments.

**Figure 2:**
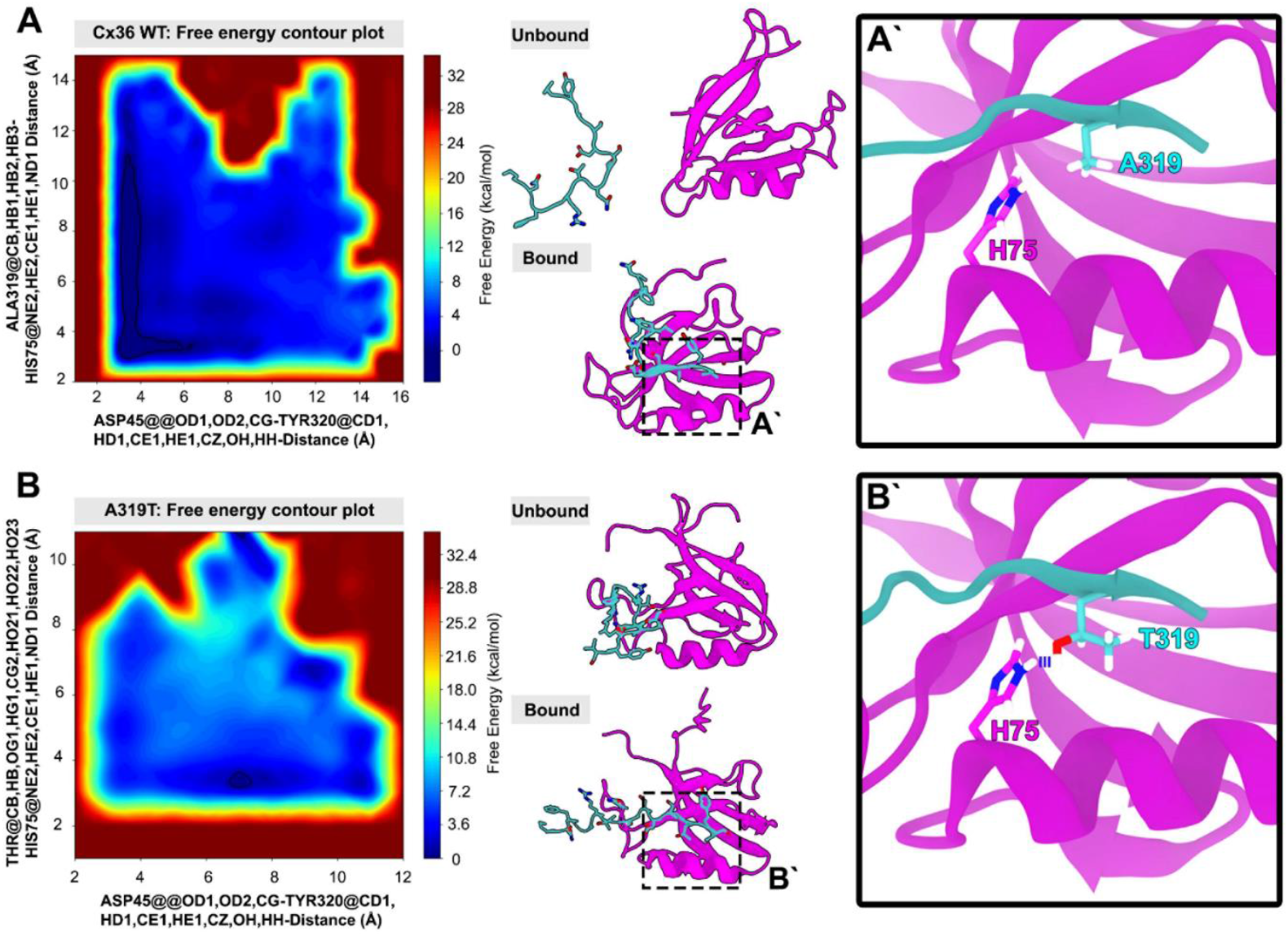
GaMD free energy landscapes of WT and A319T Cx36 PBM in complex with PDZ1. (A) Free energy contour plot of the WT PBM–PDZ1 complex show two basins corresponding to bound and unbound states. The bound basin is deep but broad, consistent with an unstable interaction. Middle panel: representative structures of unbound and bound states. Right panel (A′): close-up of the bound state showing residue A319 positioned to allow a stable hydrogen bond between the PBM backbone and H75 of PDZ1. (B) Free energy contour plot of the A319T variant shows a deeper and narrower bound basin, consistent with enhanced stability of the PDZ1–PBM complex. Middle panel: representative unbound and bound conformations. Right panel (B′): the threonine substitution provides an additional hydroxyl group that forms a more favorable hydrogen-bonding geometry with H75, reinforcing the anchoring interaction and stabilizing the bound state compared to WT.

### Amino acids adjacent to the canonical PDZ binding motif in Cx36 determine PDZ1 binding

In the next set of experiments, we searched for additional residues that Cx36 has evolved to sustain low affinity interactions with ZO-1. Our group has previously shown that phosphorylation of S315, an important CAMKII target three amino acids upstream from the PBM, is sufficient to prevent binding of the 10^th^ PDZ domain of the multi PDZ domain protein 1 (MUPP1) (40). This study demonstrated that residues that are located outside the canonical PBM in Cx36 determine the stability of PDZ interactions. Since we only addressed the role of a single posttranslational modification of a residue that is considerably far away, we wondered if amino acids that are directly adjacent to the PBM are evolutionarily tuned to support low affinity interactions as well. A sequence alignment of GJD2 and GJD1 C-termini from various species revealed that both genes carry acidic amino acids at position -4 (Figure 3A, labeled in blue) that are highly conserved in all tested vertebrates (Figure 3A, aspartate (D) for GJD2 and glutamate (E) for GJD1). To test the functional relevance of these negative charges, we substituted aspartate 317 (D317) in Cx36 with amino acids that have different chemical properties: alanine (A), arginine (R), glutamate (E) and phenylalanine (F). We co-transfected each of these mutants with Venus-ZO1-PDZ1 in HEK293T cells and performed GFP-Trap pull downs to assess PDZ binding. Out of all Cx36 variants that were tested, we observed the weakest interaction for the wild type whereas each of the D317 mutants caused a substantial increase in binding (Figure 3B). Especially the hydrophobic mutations D317A and D317F, each of which increased binding by a factor of 5-7 times, showed the most pronounced effects. These results are in line with a previous study by Zhang et al. (41) who reported that ideal PDZ binding motifs for PDZ1 (ZO-1) (ideal in terms of affinity) exhibit a preference for hydrophobic amino acids at position -4. Moreover, the finding that negative charges as the “default” at position 317 result in the weakest interactions, while amino acids with basic or hydrophobic side chains significantly enhance binding, indicates that D317 is evolutionarily tuned to support low affinity interactions. To test if this rule also applies to the closely related GJD1 protein Cx34.1 in zebrafish and the ZO-1 orthologue TJP1b, we generated a similar set of Cx34.1 mutants and repeated the binding assay (Figure 3B). In these experiments, we only saw a small, statistically insignificant increase for the E295A construct, while the E295F mutant doubled PDZ1 binding. Curiously, E295D, which converts the Cx34.1 C-terminus to the GJD2 sequence, produced a small, but statistically insignificant decrease in binding. Additionally, we tested binding of each Cx36 D317 mutant to the first PDZ domain of ZO-2. In these experiments we observed a similar preference for hydrophobic amino acids although the effects were less pronounced, with D317A and D317F causing a 2-4 times increase in binding.

**Figure 3:**
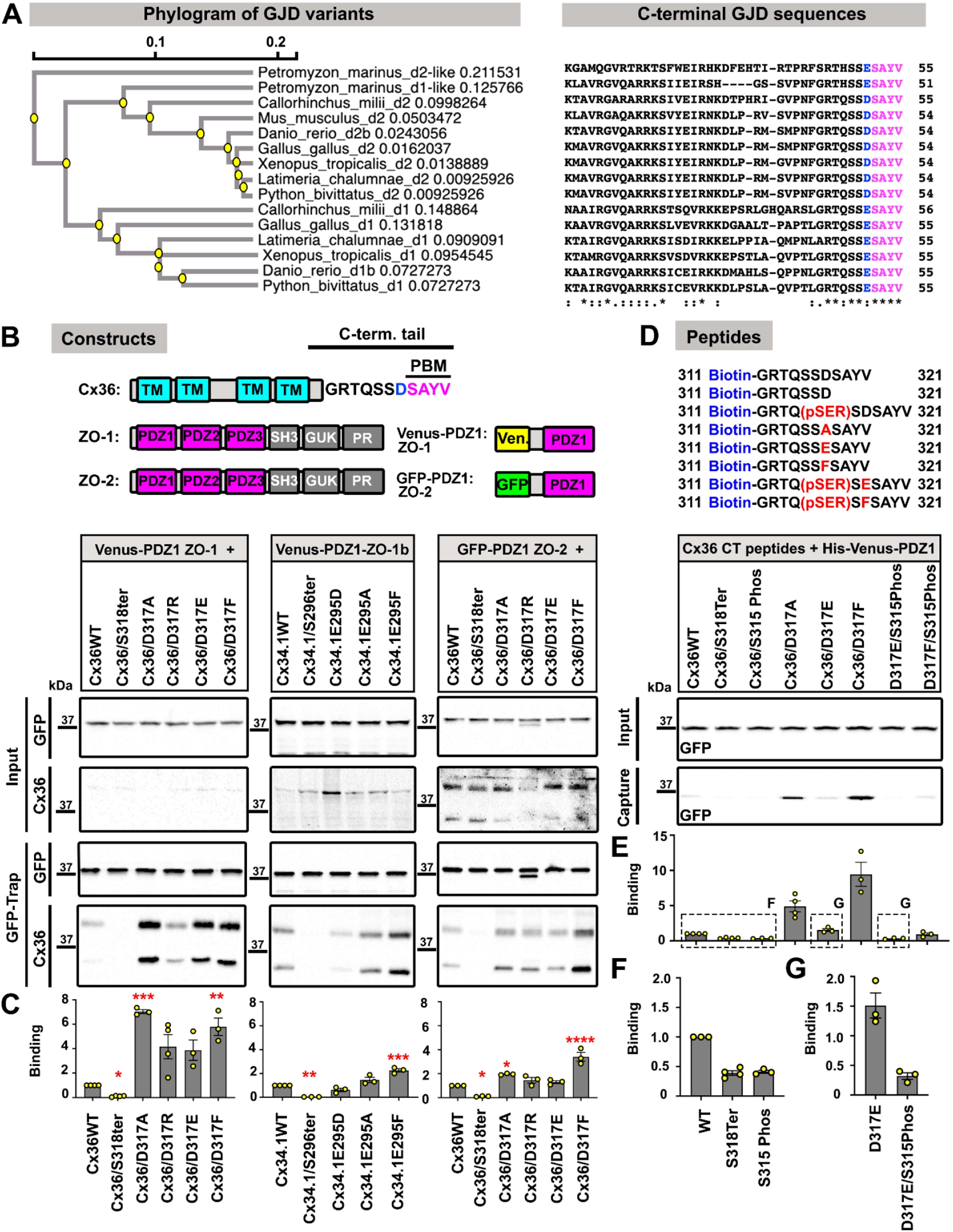
Aspartate 317, an amino acid outside the canonical PDZ binding motif of Cx36, reduces ZO-1 binding. **(A)** Phylogenetic tree depicting the evolutionary relationship of the GJD genes from various species. Sequence alignment of the GJD C-termini from various species. The sequence of the PDZ binding motif SAYV and the acidic amino acids aspartate and glutamate are highly conserved in the GJD1 and GJD2 genes from all tested species. **(B)** Domain organization of constructs used in this experiment. Aspartate 317 in Cx36 is highlighted in blue. This residue was mutated to amino acids with different chemical properties and their effect on PDZ1-ZO-1 binding was tested using GFP Trap. Cx36/Cx34.1 aspartate/glutamate mutants were co-expressed with the Venus-PDZ-1 (*Homo sapiens*) or Venus-PDZ1-ZO-1b (*Danio rerio*) construct in HEK293T and binding was assessed via Co-IP after cell lysis. **(C)** Densitometric quantification of Cx36/Cx34.1 mutants binding normalized to the wild-type variant. Significance was determined using a one-way ANOVA. Cx36 and PDZ1 (ZO-1): WT vs. D317A, p < 0.0001; WT vs. D317F, p < 0.0003. Cx34.1 and PDZ1 (ZO-1-b): WT vs. E295F, p < 0.0002. Cx36 and PDZ1 (ZO-2): WT vs. D317A, p < 0.0137; WT vs. D317F, p < 0.0001. **(D)** Sequence of biotinylated peptides used in this experiment. These peptides correspond to the individual D317 mutants used in A, except for the S315 phosphorylated peptides. All peptides were loaded onto magnetic streptavidin beads and binding was tested using His-Venus-PDZ1 (ZO-1). In line with the Co-IP data, D317A and D317F peptides showed a marked increase in PDZ1 binding. Serine 315 phosphorylated peptides show reduced binding in comparison to non-phosphorylated peptides. **(F)** Densitometric quantification of Cx36 in peptide pull-downs normalized to the wild-type pull-down. Each pull-down was replicated at least three times. **(F-G)** Binding of the phosphorylated peptides illustrated in D plotted in a magnified scale.

To complement our IP experiments, we tested if the effects we observed for the D317 variants can be replicated with peptide pull-downs. Each D317 peptide (peptide sequences are shown in Figure 3D) was coated onto streptavidin beads and probed with purified His-Venus-PDZ1 (ZO-1) to assess in binding. Overall, the peptide pull-downs confirmed the effects we have seen in CO-IP experiments. Each of the hydrophobic D317 substitutions D317A and D317F, caused the most substantial changes and increased binding by a factor 5 and 10 respectively. In addition to the D317 peptide we also tested the effect of serine 315 phosphorylation. Consistent with our previous reports (40), we observed that phosphorylation of Serine 315 is sufficient to eliminate PDZ1 binding entirely (Figure 3E-G; Note that the S315 phos and 318ter show the same yield in PDZ1, which mostly represents background). Interestingly, when we performed the pull-down with a phosphorylated version of the D317F peptide, we saw a clear reduction in binding despite the enhancing effect of phenylalanine at position 317 (Figure 3D and E). In contrast to the phosphorylated wild-type peptide, however, some binding still remained, suggesting that phosphorylation of serine 315 may be insufficient to disrupt stronger interactions entirely. Thus, one possible reason for the evolution of low affinity interactions between Cx36 and ZO-1 may lie in their flexibility and sensitivity to posttranslational modifications that allow the scaffold to dissociate from the channel under certain conditions.

### Functional Characterization of A319T mutants in transfected HeLa cells

To understand how enhancing the ZO-1/Cx36 interaction might affect the ability of Cx36 to assemble into gap junctions, we transfected the A319T mutant and wildtype Cx36 into HeLa cells and compared the mean size of plaques that were formed by both variants in µm^3^. For each region of interest, we acquired a sufficient number of slices within a z-stack (z-spacing=240nm) to ensure that the gap junction in its entire size was imaged. The size of each gap junction was determined using the *3d simple segmentation* and *measure3D* function in *ImageJ* as previously described (40, 42). This analysis revealed no significant difference in the mean size of gap junctions for each condition (Figure 4A and B). According to our analysis, both variants formed gap junctions with a mean volume of around 15 µm^3^. In addition to size, we also compared the mobility of both Cx36 variants within the gap junction plaque using fluorescence recovery after photobleaching (FRAP). To image the connexin in a FRAP paradigm, we generated a construct with an internal YFP tag in the C-terminus that exposes the 19 C-terminal amino acids of Cx36 to ensure functional PDZ interactions (1, 23). For each condition we bleached a small region within the gap junction and monitored the recovery for 100s after the bleaching step (Figure 4C). We calculated the recovery constant tau for each condition. In comparison to wild type Cx36, the A319T mutant exhibited slower recovery and a lower recovery constant (Figure 4D). This difference was insignificant, however.

**Figure 4:**
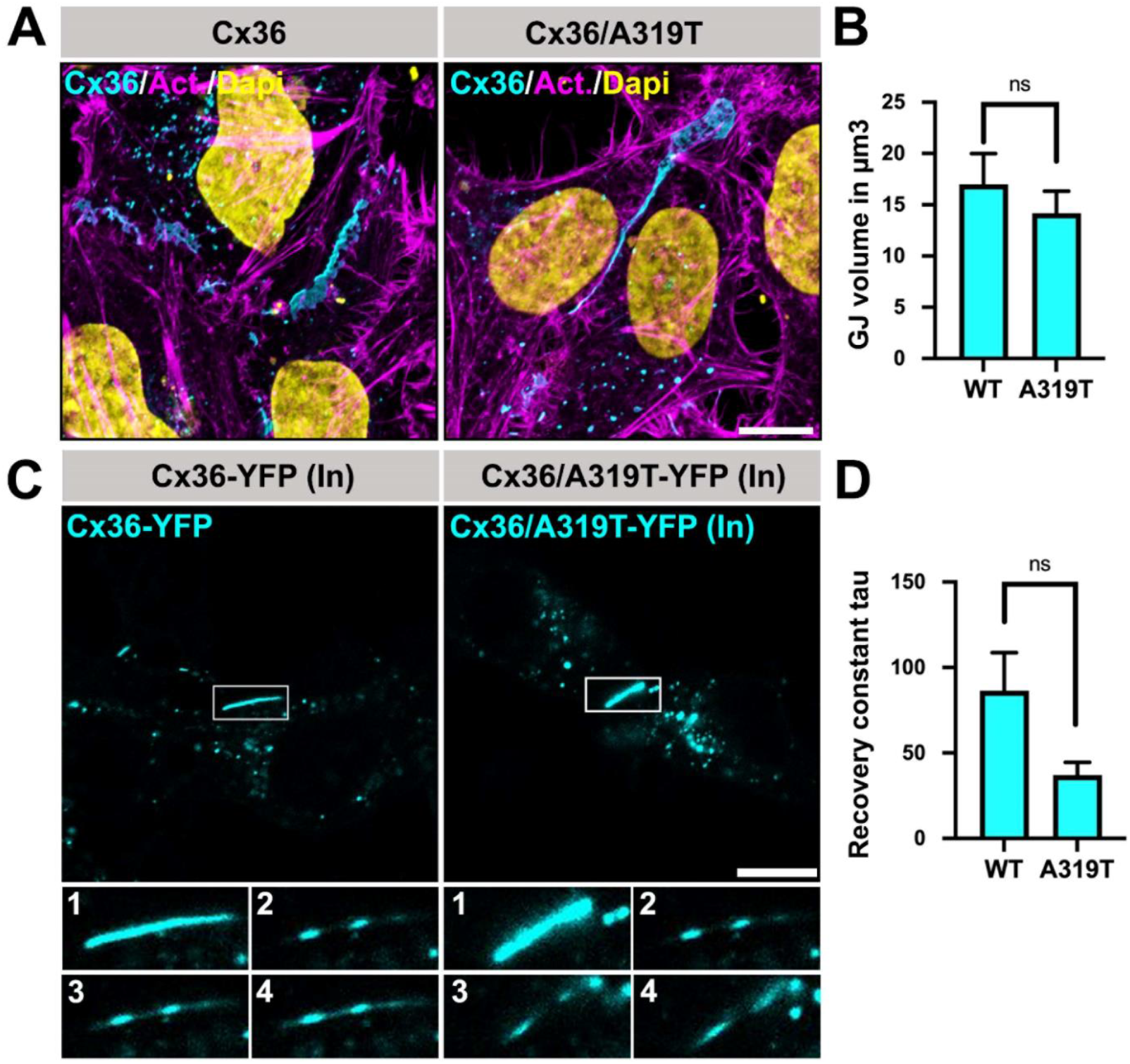
Functional characterization of the Cx36/A319T mutant in HeLa and N2A cells. **(A-B)** Double labeling of actin filaments with phalloidin and Cx36 in transfected HeLa cells. Actin filaments are associated with the periphery of each gap junction. Gap junctions of Cx36 and A319T transfected cells don’t differ significantly in size (Data are presented as mean with standard error of mean, p-value: 0.3854) Scale: 10 µm. **(C)** Frap experiments comparing the mobility of Cx36-YFP (In) and Cx36/A319T-YFP (In). Magnified insets 1-4 show the region of interest (ROI) within the gap junction for each time point in the frap paradigm: 1: ROI before bleach, 2: Bleached ROI, 3: 50 seconds post bleach, 4: 100 seconds post bleach. (D) Plot with mean recovery constant tau. The recovery constant is lower for the A319T mutant suggesting reduced mobility. Mann-Whitney U test indicates that these differences are insignificant. Data are presented as mean with standard error of mean, p-value: 0.0796.

## Discussion

In the present study, we combined enhanced-sampling molecular methods and pull-down assays to identify residues in the C-terminus of Cx36 that determine its weak interaction with the first PDZ domain of ZO-1 (Figure 5 A-C). The first critical amino acid we describe is alanine 319. Our GaMD simulations indicate that this residue is rather mobile and unable to adopt a fixed position within the binding pocket of PDZ1, which could be a contributing factor to the weak and transient nature of the Cx36/PDZ1 interaction. This argument is supported by the massive increase in binding we observed when we substituted this amino acid with threonine. According to our structural model, this increase is caused by a hydrogen bond between threonine 319 and histidine 75 in PDZ1 of ZO-1, allowing threonine to occupy a stable position. The function of this histidine as a hydrogen bond donor is conserved in many PDZ proteins (43), which explains why we observed the same effects for ZO-1 and ZO-2. Interestingly, the A319T substitution not only introduces an additional hydrogen bond, but also converts the PBM of Cx36 from a class II PDZ motif, which is characterized by two hydrophobic amino acids creating the motif Φ-X-Φ, to a class I motif S/T-X-Φ, containing a serine or a threonine, next to a random and a hydrophobic amino acid at the C-terminus. Despite their distinct chemical signatures, both motifs were shown to be compatible with the first PDZ domain of ZO-1 (39, 44). This promiscuity for different classes of PDZ motifs does not apply to all PDZ domains and appears to reflect the broad range of biological functions ZO-1 has to accomplish, considering that it functions as a major scaffold for a variety of different cell junctions including gap junctions, tight junctions and adherens junctions (26, 27, 39, 45).

**Figure 5:**
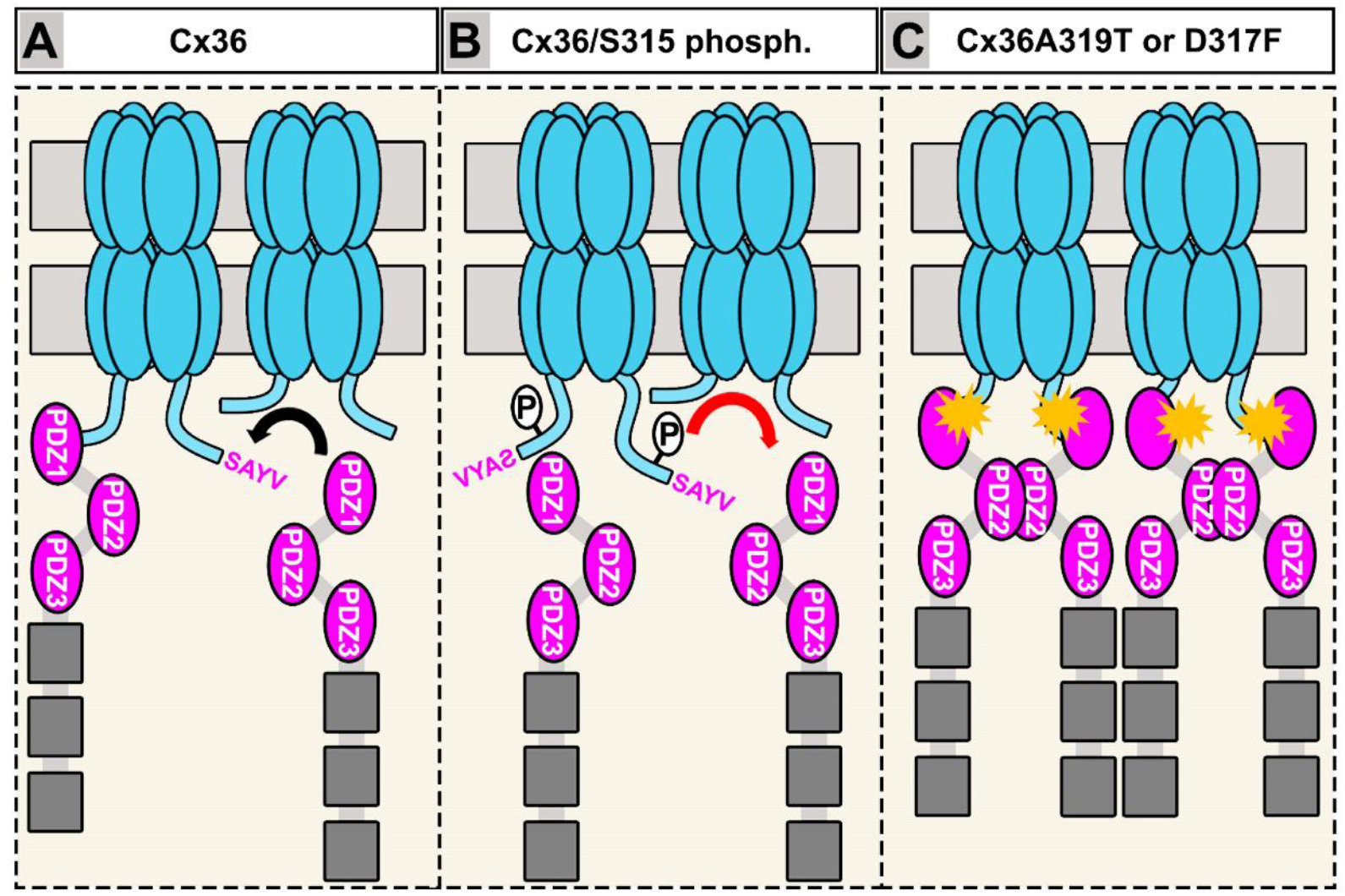
Mechanisms affecting ZO-1 binding to Cx36. **(A-B)** Binding of Cx36 to ZO-1 is weak and can be regulated by phosphorylation of S315 which might cause PDZ1 to dissociate from the channel. **(C)** The A319T and the D317F mutation we have tested in this study massively increase binding of ZO-1 to Cx36.

### Could ZO-1 potentially impact gating of Cx36 channels?

Consistent with our previous study (40), we have demonstrated that phosphorylation of serine 315 in the C-terminus of Cx36 prevents PDZ1 binding. Based on the evidence presented in this study, we propose that Cx36 has evolved the ability to form low affinity complexes with ZO-1 in order to remain flexible and sensitive to activity dependent modifications such as phosphorylation, allowing the channel to enter different functional states. This hypothesis is supported by the observation that phosphorylation of the D317F mutant failed to dissociate it fully from PDZ1, even though binding was significantly reduced in comparison to the non-phosphorylated peptide. As it is well established that phosphorylation of Cx36 potentiates electrical coupling (18, 19, 46), our findings raise the question why it also results in ZO-1 unbinding? Is it necessary to release Cx36 from ZO-1 in order to open the channel? Although structural models have been published (47, 48), it is entirely unclear how phosphorylation enables the connexin to enter a functional state in which its conductance is increased. With respect to ion channel function, scaffolding proteins such as ZO-1 or PSD-95 are mainly described as platforms that allow the assembly of signaling complexes into nano domains (49, 50). Examples in which scaffolding proteins function as sort of allosteric regulator controlling channel gating are rare (51). However, it is conceivable that ZO-1 binding imposes some sort of conformational change that locks the channel in a closed state. This hypothesis would for instance explain why phosphorylation of serine 315 is necessary to increase electrical coupling. Moreover, for Cx36 containing gap junctions in cerebellar Golgi cells it has been shown that only 18% all channels within a plaque are actually open (52). It is tempting to speculate that these 18% represent the fraction of channels that have dissociated from ZO-1. However, to fully understand the impact of ZO-1 on channel conductance, detailed electrophysiological experiments and insights into stoichiometry of these complexes are necessary.

### Could enhanced ZO-1 binding limit the mobility of Cx36 channels within the gap junction?

While our study identified different ways to manipulate ZO-1 binding, we currently lack an appropriate *in vivo* model that would allow us to assess defects in electrical synapse function arising from enhanced PDZ interactions, which is why we decided to study the effects of the A319T mutation in a simple expression system. These experiments, however, revealed no obvious changes in gap junction size or the mobility of channels. Although the differences in mobility between wild type Cx36 and the A319T mutant appeared to be insignificant, we noticed a visible trend towards lower recovery constants for the A319T condition, which may result from increased ZO-1 binding limiting the lateral diffusion of channels within the gap junction plaque. Moreover, the lack of an “obvious” phenotype in our FRAP experiment may be due to the variability that is introduced by transient transfections. Within a cell culture dish, each cell takes up varying amounts of the vector resulting in considerable differences in the expression level of the fusion protein. This variation might make it more difficult to detect consistent effects on the recovery constant tau.

### Is there a connection between endocytosis of Cx36 and ZO-1 binding?

Previously, Thevenin et al., (53) have demonstrated that phospho-dead S373A Cx43 mutants in transfected Madin-Darby canine kidney (MDCK) cells show increased ZO-1 binding, abnormally large gap junctions and prolonged lifetimes. As we have described for Cx36, the interaction between Cx43 and ZO-1 is regulated by phosphorylation of S373, which serves as a binding switch that regulates the size of gap junctions under hypoxic conditions (54). Lack of phosphorylation in S373A mutants consequently increases the association with ZO-1 and results in the described phenotypes (53). However, it is not entirely clear how these effects are connected and why enhanced ZO-1 binding causes abnormal gap junction growth. A possible link between these two effects, as outlined by Thevenin et al., could be limited access of endocytosis proteins that need to engage in an intimate interaction with Cx43 in order to initiate the internalization of gap junction channels (55). A tight interaction with large scaffolds such as ZO-1 could potentially block binding of ubiquitin ligases or ubiquitin receptors such as EPS15, which would ultimately inhibit endocytosis and result in abnormal gap junction growth. This hypothesis also fits well with the characteristic “perinexus” distribution of ZO-1, in which Cx43 channels in the periphery of the plaque interact with the scaffold, while the center of the gap junction, from which older channels are usually removed, lack ZO-1, suggesting that disengagement from ZO-1 is necessary for functional endocytosis (56, 57). A similar mechanism might apply to Cx36 (and synapses that are formed by the GJD orthologues in zebrafish) as it was shown that intradendritic application of peptides that mimic the C-terminus of Cx35 (and the PBM) result in increased internalization of Cx35 in Mauthner cells, suggesting that ZO-1 has a stabilizing effect opposing endocytosis. Moreover, we have previously reported that endocytosis proteins such as epidermal growth factor receptor pathway substrate 15 like 1 (Eps15l1) colocalize with more than 70% of all Cx36 puncta in the inner retina (2). This observation is consistent with the fast turnover rate of Cx36 (23, 24) and suggests that the connexin content of an electrical synapse is tightly controlled. The weak binding affinity for ZO-1 could be useful in this context and might allow quick rearrangements of the scaffold to facilitate interactions of Cx36 with endocytosis proteins.

## Material and Methods

### Cell Culture

Human embryonic kidney 293 T cells (HEK293T/17; catalog #CRL-11268; ATCC, Manassas, VA, USA) were cultivated in Dulbecco’s Modified Eagle Medium (DMEM) supplemented with 10% fetal bovine serum (FBS), 1% penicillin and streptomycin, and 1% non-essential amino acids (all Thermo Fisher Scientific, Rockford, IL, USA) at 37 °C in a humidified atmosphere with 5% CO2. HeLa cells (catalog #CCL2, ATC) were cultivated under similar conditions. Instead of DMEM these cells were grown in MEM.

### Co-IP experiments

Immunoprecipitation experiments were carried out as previously described (1, 2). 1.5 ×10^6^ HEK293T cells were plated onto 60mm dishes and grown overnight. At the next day, cells were co-transfected with 2µg of the Cx36 pcDNA vector or specific Cx36 mutants and 2 µg of Venus-PDZ1 (ZO-1) expression vector (1) using Geneporter 2 (Amsbio). 24h post-transfections, cells were removed from the dish and pelleted (centrifugation for 5 min at 5000g at RT) in phosphate buffered saline (PBS) (pH 7.4). The cell pellets were resuspended and lysed in immunoprecipitation (IP) buffer containing 200mM NaCl, 2mM EDTA, 1% Triton X-100, 50mM Tris HCL (pH 7.5) supplemented with protease inhibitors (Roche) and 1mM DTT. The lysate was chilled on ice for 30 min and sonicated 3x for 30 seconds every 10 min. Afterwards, the sample was centrifuged at 10,000 g for 10 min and the resulting supernatant was incubated with 20µl ChromoTek GFP-Trap® agarose beads (gta, Proteintech) overnight at 4ºC on a rotating platform. The next day, IP samples were centrifuged at 2500 g for 5 min and washed three times in IP buffer. Bound proteins were eluted with 60 µl of 1x Laemmli buffer (BioRad, 1610747) for 5 min at 95ºC.

### Purification of His-Venus-PDZ1 (ZO-1)

HEK293T cells were plated onto 100mm dishes and grown to 80% confluency. 18 dishes of confluent cells were transfected with 10µg His-Venus-PDZ1 each using lipofectamine 3000 (catalog#L3000001, Thermofisher scientific). 48h post transfection cells were collected and lysed in IP buffer. Cell lysates were centrifuged at 10.000g for 10 minutes. The supernatant was applied to a Nickel column and incubated for 1h on a rotating platform at 4º C. Afterwards, the supernatant was allowed to flow through, and the column was washed 3x with wash buffer (50mM Tris pH 8.0, 300mM NaCl and 30mM Imidazole). Bound proteins were eluted using elution buffer containing 50mM Tris (pH 8.0), 50mM NaCl and 300mM Imidazole. The elution buffer was substituted with binding buffer (10 mM KH_2_PO_4_, 150mM NaCl, 3mM KCl and 0.1% pluronic acid, pH 7.4) using a PD10 column (cat#17085101, Cytiva).

### Peptide pull downs

Biotinylated peptides corresponding to the C-terminal 12 amino acids of Cx36 and individual mutant variants were synthesized by Genscript (peptide sequences shown in Table 1). Each peptide was solved in DMSO with a concentration of 20mM.

**Table 1.** Sequence of C-terminal Cx36 peptides used in this study.

| C-terminal peptide | Sequence |
| --- | --- |
| Cx36 | Biotin-GRTQSSDSAYV |
| Cx36/S318Ter | Biotin-GRTQSSD |
| Cx36/A319T | Biotin-GRTQSSDSTYV |
| Cx36/S315 Phos | Biotin-GRTQS(P)SDSAYV |
| Cx36/D317A | Biotin-GRTQSSASAYV |
| Cx36/D317E | Biotin-GRTQSSESAYV |
| Cx36/D317F | Biotin-GRTQSSFSAYV |
| Cx36/D317E/S315Phos | Biotin-GRTQS(P)SESAYV |
| Cx36/D317E/S315Phos | Biotin-GRTQS(P)SFSAYV |

For each pull-down experiment, 1 µl of each peptide was combined with 50 µl Streptavidin C1 Dynabeads™, MyOne™ (catalog#65001, Thermofisher) and 900 µl binding buffer and incubated for 30 minutes at RT on a rotating platform. Afterwards, the beads were washed 3x with binding buffer and combined with 180 µl of the purified PDZ domain and 270 µl binding buffer. The sample was incubated for 1h at 4ºC on a rotating platform and washed afterwards three times with binding buffer. Bound proteins were eluted with 1x Laemmli buffer (BioRad, catlog#1610747) for 5 min at 95ºC.

### Western blot

Western blots were carried as described previously (2). Protein samples were separated via SDS-PAGE containing 10% polyacrylamide. Afterwards, proteins were transferred to nitrocellulose membrane using the transblot turbo system (Catlog#1704150, BioRad). Nitrocellulose membranes were blocked with 5% dried milk powder in TBST (20 mM Tris pH 7.5, 150 mM NaCl, 0.2% Tween 20) for 30 min at RT and incubated overnight in the primary antibodies diluted in blocking solution at 4ºC on a rotating platform. The following primary antibodies were used: Cx36, 1:500 (Clone: 8F6, catalog#MAB3045, Millipore); Cx36, 1:500 (Clone: 1E5H5, catalog#37-4600, Thermofisher), GFP, 1:1000 (#2956, Cell Signaling) and ZO-1, 1:500 (Clone: ZO1-1A12, catalog#33-9100, Thermofisher). At the next day, blotting membranes were washed 3×10 min in TBST and incubated with secondary antibodies. The following secondary antibodies were used: goat anti mouse HRP, 1:1000 (34130, Thermofisher) and goat anti rabbit, 1:1000 (34160, Thermofisher). Afterwards the membranes were washed 3×10 min in TBST and imaged using enhance chemiluminescence as described before (2).

### Immunocytochemistry

Coverslips with HEK293T and HeLa cells were fixed with 2% paraformaldehyde solved in PBS, pH 7.4 for 15 min at RT. Afterwards coverslips were washed 3×10 min in PBS and incubated in the primary antibody solution containing 10% normal donkey serum in 0.5% Triton X-100 in PBS overnight at 4ºC. The following primary antibodies were used: Cx36, 1:500 (Clone: 8F6, catalog#MAB3045, Millipore), Cx36, 1:250 (catalog#36-4600, Thermofisher) and ZO-1, 1:250 (Clone: ZO1-1A12, catalog#33-9100, Thermofisher). The next day the cover slips were washed 3×10 min in PBS and incubated in the secondary antibody solution containing 10% normal donkey serum in 0.5% Triton X-100 in PBS for 1h at RT under light protection. The following secondary antibodies were used: Donkey anti rabbit Cy3, catalog#711-165-152, 1:250 and donkey anti mouse 488, catalog#711-545-151, 1:250, Jackson ImmunoResearch. Afterwards, the coverslips were washed 3×10 min in PBS and mounted with Vectashield (catalog#UX-93952-24, Vector Laboratories).

### Image acquisition and volume quantification

The volume and frequency of Cx36 gap junctions in transfected HeLa cells was measured as previously described (37, 40, 42). Cx36 expressing cell clusters were scanned using a 60x oil objective (Zeiss LSM 800) (1024×1024 format, 0.99 nm x 0.99 nm pixel size) with a stack size (ranging from 18 to 45 planes depending on the size of each gap junction; z-spacing: 240nm) containing a sufficient number of planes to cover the entire volume of individual gap junctions. The volume of each junction we captured was quantified using the *3D Manager* plugin in Fiji (58). First, images were thresholded using the Otsu threshold and then segmented using the 3D simple segmentation function. Individual regions of interests (ROI)/gap junctions were selected with the *3D Manager* and measured using the *measure 3D* function.

The frequence of gap junctions per cell was measured using the *cell counter* plugin. For each cell cluster, confocal scans were maximum projected (20 planes per cluster). In these projections, the number of gap junctions in each cluster was quantified using the cell counter plugin. The number of gap junctions was divided by number of cell pairs.

### Molecular Modeling and Molecular Dynamics Simulations

#### Experimental considerations

We initially employed endpoint free energy methods (MMGBSA/MMPBSA) implemented in both GROMACS and AMBER to compare binding energetics between the WT and variant complexes. However, these calculations yielded very similar ΔG values for both systems, with differences falling within the expected statistical error of the method. This result indicated that standard MMGBSA, while computationally efficient, relies on static snapshots and averages over limited conformational sampling, often neglecting entropic contributions and dynamic rearrangements. As a result, it can fail to capture subtle yet biologically relevant binding differences; particularly in systems where the thermodynamics are shaped by conformational heterogeneity or metastable microstates. In our case, although the WT and variant complexes displayed comparable net binding enthalpy, they differed markedly in the organization and dynamics of hydrogen bonds, electrostatic contacts, and hydrophobic networks throughout the trajectory. These structural and energetic redistributions were not reflected in MMGBSA-derived ΔG values, rendering the method insensitive to experimentally supported differences in binding behavior. Similar limitations have been reported in other protein–peptide systems, where MMGBSA failed to distinguish between ligands with distinct affinities, while enhanced sampling approaches like GaMD, metadynamics, or weighted-ensemble simulations succeeded by incorporating entropy and capturing multiple binding modes (59-61). To address this, we applied Ligand3 (62), a trajectory reweighting strategy that accounts for microstate-specific contributions to free energy and enables a more nuanced analysis of binding energetics.

#### System Preparation and Equilibration

All molecular dynamics simulations were performed using the AMBER 24.4 software suite (63). Initial atomic models of the Connexin 36/ZO-1 complexes were generated using AlphaFold-Multimer and subsequently prepared for simulation. The Connexin 36/ZO-1 protein complex was first immersed in an explicit solvent environment. A rectangular periodic box of TIP3P water molecules was built around the complex, with at least a 10 Å buffer separating the protein from the box edges. Sodium and chloride ions were added to neutralize the system and approximate a physiological salt concentration (0.15 M). Protein parameters were assigned from the AMBER all-atom force field (ff14SB or equivalent), and Lennard-Jones and partial charge parameters for ions and water were taken from standard AMBER libraries (TIP3P water model).

Prior to production simulations, the solvated system was energy-minimized and equilibrated. Initial energy minimization relieved any steric clashes, typically using a combination of steepest descent and conjugate gradient algorithms under harmonic restraints on the protein heavy atoms. The system was then gradually heated from 0 to 300 K under constant volume (NVT) dynamics, using a Langevin thermostat (collision frequency of 2 ps^−1^) to control temperature. During heating, a positional restraint of 5 kcal·mol^−1^·Å^−2^ was applied to protein heavy atoms to maintain the complex geometry. Next, equilibrium was achieved under isothermal-isobaric conditions (NPT) at 1 atm and 300 K, employing a Monte Carlo barostat for pressure control. The positional restraints were gradually reduced and finally removed over a series of short simulation stages, allowing the protein and solvent to relax fully. All bonds involving hydrogen atoms were constrained with the SHAKE algorithm, enabling a 2 fs integration time step throughout the simulations.

#### GaMD Production Simulations and Reweighting

Production simulations were carried out using Gaussian accelerated molecular dynamics (GaMD) (64). GaMD is an enhanced sampling approach in which a small harmonic boost potential is added to the system’s potential energy to smooth the energy landscape and reduce energy barriers. In GaMD, the magnitude of the applied boost is governed by a user-defined threshold energy and is designed such that the distribution of the boost potential is approximately Gaussian. This allows for rigorous energetic reweighting via a cumulant expansion (the “Gaussian approximation”) to recover accurate unbiased free energy landscapes from the simulation (65). For our GaMD implementation in AMBER, we employed the LiGaMD3 (62) dual-boost scheme, applying one boost potential to the dihedral (torsional) component of the potential energy and a second boost to the total potential energy of the system.

GaMD boost parameters were chosen according to published guidelines to ensure both sampling and reweighting. In particular, the threshold energies for each boost were set such that the standard deviation of the boost potential (σ_ΔV_) did not exceed ∼6.0 kcal/mol (roughly 10 k_BT at 300 K). This criterion is followed to maintain a narrow, near-Gaussian distribution of the boost potential, as recommended for accurate reweighting (65). During an initial calibration phase, short GaMD trials were performed to adjust the boost parameters (e.g. the σ0 values for dihedral and total boosts) until both boost distributions met the GaMD stability criteria (typically, anharmonicity <0.3 and boost mean<ΔE_max). Once parameters were finalized, full production GaMD simulations were launched.

At least 25 independent GaMD production runs (replicates) were conducted, each 500 ns in length, yielding an aggregate simulation time of 15 μs or more. Each replicate simulation was initiated from the equilibrated structure of the Cx36/ZO-1 complex, with uniquely randomized initial velocities to ensure independent trajectories. By distributing the sampling across multiple long replicates, we increased the diversity of explored conformational space and the likelihood of observing rare unbinding events. GaMD (including its specialized variants) has been demonstrated to capture repetitive binding and dissociation events in protein–ligand systems, providing reliable estimates of binding thermodynamics and kinetics (64). Following this best-practice approach, the extensive GaMD sampling in our study was designed to thoroughly explore the Connexin 36/ZO-1 interaction landscape and improve the statistical confidence of the results.

To obtain unbiased free energy surfaces from the GaMD simulations, we performed post-simulation reweighting of the trajectories using the PyReweighting toolkit (65). PyReweighting is a Python-based toolkit that implements several statistical reweighting methods for accelerated MD simulations. In this work, we utilized the cumulant expansion to second order (Gaussian approximation) to reweight the GaMD trajectories. For each saved trajectory frame, the instantaneous boost potential ΔV applied by GaMD was recorded. The ensemble-averaged Boltzmann factor of the boost was then computed via the second-order cumulant expansion, which uses the first and second cumulants of the ΔV distribution (i.e. the mean boost and its variance) to approximate the reweighting factor. This procedure corrects for the bias introduced by the GaMD boosts and yields a weight for each frame that reflects its probability in the canonical (unbiased) ensemble. Frames were thus reweighted to reconstruct the original potential of mean force (PMF) for the system.

Using the PyReweighting toolkit, we calculated free energy profiles along relevant reaction coordinates describing the Cx36/ZO-1 binding interface. In particular, one- and two-dimensional PMF analyses were carried out by binning the GaMD trajectory data in chosen coordinate space and applying the corresponding frame weights. For example, distance-based reaction coordinates (such as the inter-protein center-of-mass distance or specific contact distances) and orientation/orientational order parameters were considered to characterize the binding interaction. The free energy F as a function of a given coordinate was obtained from the reweighted probability P using *F(x) = -k*_*B*_*T ln P(x)*. To ensure sufficient sampling in all regions of the free energy surface, we excluded any grid bins that contained too few samples. Bins with fewer than a minimum number of frames (determined by gradually increasing the cutoff until the resulting PMF converged) were omitted from the PMF construction. The final reweighted free energy surfaces provide an unbiased representation of the binding energetics and conformational landscape of the Connexin 36/ZO-1 complex, as derived from our enhanced sampling simulations.

#### Sequence analysis

*Gjd1* and *Gjd2* genes in a variety of vertebrate species were discovered in whole genome sequence data using TBlastN with perch Cx35.1 or zebrafish Cx34.1 predicted amino acid sequences as the query. Sources of sequences included in the comparison are shown in Table 2. Predicted amino acid sequences of the C-terminal domain beginning after the fourth transmembrane domain were aligned with Clustal Omega (66), version 1.2.4 using the EMBL Clustal server (accessed at https://www.ebi.ac.uk/jdispatcher/msa/clustalo).

**Table 2.** Sources of sequences used for Gjd1/Gjd2 alignment.

| Species | Gene | Chromosome | Annotated gene name | Accession |
| --- | --- | --- | --- | --- |
| *Petromyzon_marinus* | Gjd1-like | 33 | LOC116948513 | NC_133699 |
|  | Gjd2-like | 6 | LOC116939521 | NC_133672 |
| *Callorhinchus milii* | Gjd1 | unplaced | LOC103173305 | XM_042345617 |
|  | Gjd2 | unplaced | gjd2b | XM_007888024 |
| *Danio_rerio* | gjd1b | 15 | LOC100331216 | XM_009291479 |
|  | gjd2b | 20 | gjd2b | NM_194420 |
| *Latimeria_chalumnae* | Gjd1 | 29 | LOC102366134 | XM_005987869 |
|  | Gjd2 | 11 | GJD2B | XM_006004060 |
| *Xenopus_tropicalis* | Gjd1 | 8 | LOC100494762 | KAE8585750 |
|  | Gjd2 | 8 | gjd2 | XM_002932834 |
| *Mus_musculus* | Gjd2 | 2 | Gjd2 | NM_010290 |
| *Python_bivittatus* | Gjd1 | unplaced | LOC103048063 | XM_007426978 |
|  | Gjd2 | unplaced | GJD2 | XM_007434115 |
| *Gallus_gallus* | Gjd1 | 38 | LOC107049826 | XM_040655140 |
|  | Gjd2 | 5 | GJD2 | NM_204582 |

## Acknowledgements

This project was supported by the NIH grant R01EY012857 (J.O.). S.T. was funded by the *Deutsche Forschungsgemeinschaft* (DFG) (TE 1459/1-1, Walter Benjamin stipend). E.S. was supported by NIH training grant TL1TR003169 and individual grant F31EY034793. We would like to thank Dr. Adam C. Miller and Dr. Jen Michel for providing the Cx34.1 antibody and the tjp1 expression construct.

